# Conserved fiber topography of the anterior limb of the internal capsule in treatment-resistant psychiatric patients

**DOI:** 10.64898/2026.05.11.724148

**Authors:** Reem El Jammal, Hideo Suzuki, Layth S. Mattar, Thomas Hamre, Sarah Soubra, Melissa A. Ryan, Raissa K. Mathura, Sanjay J. Mathew, Anusha Allawala, Eric A. Storch, Nora Vanegas Arroyave, Garrett P. Banks, Nader Pouratian, Remi Patriat, Wayne K. Goodman, Nicole R. Provenza, Sameer A. Sheth, Eleonora Bartoli, Sarah R. Heilbronner

## Abstract

**Introduction:** The anterior limb of the internal capsule (ALIC) is a major white matter highway connecting prefrontal cortical (PFC) regions to the thalamus, brainstem, and subthalamic nucleus. Structural and functional abnormalities within the ALIC circuit have been associated with many neuropsychiatric disorders, including obsessive-compulsive disorder (OCD) and depression, and deep brain stimulation (DBS) may provide effective treatment to some of these patients. However, it remains unclear whether the well-characterized topographic organization of the ALIC observed in healthy individuals and preclinical models is preserved in treatment-resistant psychiatric populations.

**Methods:** We first used diffusion tractography to evaluate the topography of PFC and subcortical fibers through the ALIC in patients with treatment-resistant OCD (n=18) and depression (n=5). In depression patients, we also evaluated ALIC topography using cerebro-cerebral evoked potentials (CCEPs) elicited by single-pulse electrical stimulation (SPES) of DBS leads in the ALIC and recordings in the ventral PFC (vPFC).

**Results:** The topographic organization of PFC and subcortical projections is preserved in the ALIC among treatment-resistant psychiatric patients, consistent with patterns observed in healthy individuals and preclinical models. CCEP recordings in the ventral PFC showed a ventral ALIC to medial vPFC/dorsal ALIC to lateral vPFC response pattern in the left hemisphere, but not in the right.

**Conclusion:** Our findings confirm that topographic patterns within the ALIC previously identified using preclinical models and healthy controls are preserved in treatment-resistant psychiatric patients. Furthermore, by linking white matter topography to stimulation effects, this work supports more precise and individualized neuromodulatory strategies for neuropsychiatric disorders.

## Introduction

The internal capsule is a white matter highway connecting the cerebral cortex with subcortical structures. The fibers in the anterior limb of the internal capsule (ALIC) carry information specifically from the prefrontal cortex (PFC) to the thalamus, subthalamic nucleus (STN), and brainstem (Beevor and Horsley, 1890; Jbabdi et al., 2013; Safadi et al., 2018). Disruption of the ALIC and its associated circuits have been linked to a broad spectrum of neuropsychiatric symptoms (Levitt et al., 2012; Nanda et al., 2017; Mithani et al., 2020).

Although in most instances, psychiatric disorders can be effectively managed with pharmacological and behavioral therapies, stereotactic neurosurgical interventions provide a critical option for severe cases refractory to pharmacotherapy (Nuttin et al., 2014). The ALIC has emerged as a surgical target for neuromodulatory interventions in refractory neuropsychiatric disorders, likely due to its pivotal role in regulating limbic and cognitive circuits. Deep brain stimulation (DBS) of the ventral portion of the ALIC, for example, received approval from the U.S. Food and Drug Administration (FDA) under a Humanitarian Device Exemption (HDE) for treatment-resistant obsessive-compulsive disorder (OCD) (Nuttin et al., 2003; Greenberg et al., 2006; Mallet et al., 2008; Borron and Dougherty, 2022; Sheth et al., 2026). In addition, DBS of the ALIC has been investigated for other neuropsychiatric conditions, including major depressive disorder, Tourette syndrome, anorexia, and addiction, underscoring the ALIC’s broader therapeutic potential (Malone, 2010; Lozano et al., 2012; Kuhn et al., 2014; Baldermann et al., 2016; Graat et al., 2017). Beyond DBS, capsulotomy lesions also target the ALIC in OCD leading to improvements in symptoms, disability, and quality of life (Miguel et al., 2019; Cui et al., 2023; Gupta et al., 2024).

Tract-tracing in nonhuman animal models has demonstrated that the ALIC is precisely topographically organized, with PFC region of origin strongly determining where fibers travel within the bundle (Lehman et al., 2011; Coizet et al., 2017; Safadi et al., 2018). For example, dorsal PFC fibers travel dorsally within the ALIC to fibers from ventral PFC regions. ALIC fibers from the anterior dorsal PFC regions travel ventrally to fibers from the posterior dorsal PFC regions. For the ventral PFC, fibers from medial regions travel ventrally to those from lateral regions, whereas in the dorsal PFC, this pattern is reversed. This complex topography can theoretically be used to surgically target selective PFC fibers while leaving others unaffected (Jbabdi et al., 2013; Sretavan et al., 2024).

Neuroimaging tools that enable visualization and quantification of white matter connectivity can uncover the topographical organization of bundles such as the ALIC and can guide precise targeting for therapeutic interventions. Diffusion-weighted magnetic resonance imaging (dMRI) tractography provides a noninvasive tool for delineating specific white matter pathways (Le Bihan et al., 1986; Basser et al., 1994; Mori et al., 1999; Tournier et al., 2011), facilitating surgical planning aimed at modulating critical networks while reducing the risk of unwanted side effects (Kamagata et al., 2024). For example, Li and colleagues (2020) utilized dMRI tractography to determine the ideal set of connections to modify within the ALIC to achieve efficacy of DBS for OCD (Li et al., 2020). Outside of the ALIC, dMRI tractography has allowed for precise targeting of the dentato-rubro thalamic tract for essential tremor (Fenoy and Schiess, 2017; Miller et al., 2019), the subcallosal cingulate (SCC) white matter region for major depression (Riva-Posse et al., 2014; Song et al., 2023), and the STN for Parkinson’s disease (Schrock et al., 2021).

However, diffusion tractography, as the only tool we have available to noninvasively map structural connectivity, suffers from well documented inaccuracies (Thomas et al., 2014; van den Heuvel et al., 2015; Donahue et al., 2016; Schilling et al., 2019; Grier et al., 2020). It can both create spurious pathways (false positives) and fail to capture true ones (false negatives). Diffusion tractography can, however, be tailored for particular circuits to reconstruct patterns of anatomical connectivity. In order to do so, detailed understanding of the underlying anatomy must be known (Schilling et al., 2019). Notably, ALIC topography has been well-mapped through careful tract-tracing in nonhuman primates and corresponding diffusion imaging in healthy individuals (Nanda et al., 2017; Safadi et al., 2018; Sretavan et al., 2024).

We previously developed a novel data analysis pipeline that reliably reconstructs biologically validated ALIC fiber pathways from prefrontal cortical regions, strengthening the anatomical foundation for tractography in this region (Sretavan et al., 2024). This was done by applying this pipeline to MRI data from healthy control participants from the Human Connectome Project (HCP), producing 22 segmented ALIC bundles (11 per hemisphere, each representing the contribution of a unique PFC region). The pipeline reproduced established anatomical topographies across field strengths, and test-retest data revealed significantly smaller intraparticipant than interparticipant variability. In parallel, we also used HCP data to segment the ALIC according to subcortical targets (Banks et al., 2022). We were able to detect a medial-to-lateral organizational gradient, with thalamic fibers positioned medially and STN and brainstem fibers laterally. This topography, also observed in microscopy data (Axer et al., 1999), fades more anteriorly. Together, these results indicate that diffusion tractography, properly applied, can reliably reproduce established ALIC anatomy and provide a validated, patient-specific framework for tractography-guided DBS targeting and mechanistic modeling of therapeutic response.

Beyond diffusion tractography and tract-tracing, cerebro-cerebral evoked potentials (CCEPs) provide a functional connectivity map of electrical circuits. (Originally referred to as cortico-cortical evoked potentials, the frequent inclusion of electrodes positioned in white matter and subcortical targets has led to a broader definition (Lyu et al., 2025). CCEPs are elicited by delivering a small electrical potential at a specific brain region while recording short-latency activity in distributed cortical and subcortical targets. The presence and strength of CCEPs correlate with structural connectivity as estimated through diffusion tractography (Kundu et al., 2020; Crocker et al., 2021; Schmid et al., 2023). Importantly, CCEPs may be used to determine DBS target engagement, offering a direct window into how stimulation modulates circuit activity (Noor et al., 2024). Leveraging these signals can guide patient-specific adjustments in stimulation parameters of neuromodulation therapies such as DBS, thereby improving clinical efficacy, reducing side effects, and ultimately enhancing outcomes in OCD and other neuropsychiatric disorders.

It is still unclear whether the specific ALIC topographies identified in healthy individuals can be reproduced using similar diffusion tractography methods in treatment-resistant psychiatric patients. While studies focused on diffusion imaging show that abnormalities within the ALIC are linked to psychiatric disorders, these results relied on diffusion metrics rather than topographical organization (Levitt et al., 2012; Lochner et al., 2012; Safadi et al., 2018). These abnormalities and the associated circuits in psychiatric patients may indicate that topographies are disrupted or difficult to detect. Alternatively, topographies may be preserved, with connection strength or direction impacted instead. Thus, in a cohort of ALIC DBS patients (treatment-resistant OCD, and treatment-resistant depression, TRD) we set out to: 1) demonstrate that our PFC and subcortical diffusion tractography algorithm in the ALIC generates the established topographies in healthy individuals when applied to these patient cohorts; 2) evaluate the functional topographical organization of the ventral PFC (vPFC)-ALIC projection in TRD patients using a unique CCEP dataset.

## Materials and Methods

### Participants

The cohort included 18 patients with severe OCD treated with DBS (9 males, 38.77 ± 11.13 years; 9 females, 37.88 years ± 12.09 years), with pre-operative Yale-Brown Obsessive Compulsive Scale (YBOCS) (Goodman et al., 1989) scores averaging 36 ± 1.73 for males and 37.6 ± 3.35 for females. We additionally studied a cohort of 5 TRD patients (2 males, 49 ± 24.04 years; 3 females, 43 ± 12.12 years) who underwent DBS and stereoelectroencephalography (sEEG) electrode implantation to investigate electrophysiological mechanisms underlying this disorder were also included in the study. Baseline Hamilton Depression Rating Scale (HAMD) (Hamilton, 1960) scores were obtained pre-operatively twice for each subject averaging 22.5 ± 2.12 in male participants and 23 ± 3.605 in female participants. All subjects provided written informed consent to be included in the study which was approved by the Institutional Review board (IRB) at Baylor College of Medicine (BCM IRB number: H-43144 for OCD study and H-43036 for TRD study).

### Surgical procedure

All OCD patients underwent ALIC DBS electrode implantation specifically to the ventral capsule/ventral striatum (VC/VS) and/or ventral capsule/bed nucleus of the stria terminalis (VC/BNST), with 8 patients implanted with bilateral Medtronic Sensight B33015 leads and 10 patients implanted with bilateral Medtronic 3387 leads using robotic stereotactic guidance (Giridharan et al., 2022; Shofty et al., 2023). TRD patients underwent stereotactic implantation of four segmented DBS leads (Boston Scientific Cartesia) bilaterally in the VC/VS and SCC, and ten temporary sEEG recording electrodes (PMT Corporation, MN, USA) positioned using pre-operative patient-specific tractography-guided planning to help interpret network activity (Sheth et al., 2022, 2023). sEEG electrodes had 12 to 16 intracranial depth contacts each with a 0.8 mm diameter and a 3.5 center-to-center distance that enable neural activity recording in cortical, subcortical, and white matter regions. Contacts were implanted bilaterally to cover various regions of interest including the dorsolateral and dorsomedial prefrontal cortex (dlPFC, dmPFC), ventrolateral and ventromedial prefrontal cortex (vlPFC, vmPFC), medial and lateral orbitofrontal cortex (mOFC and lOFC), anterior cingulate cortex (ACC), amygdala, and middle and superior temporal gyri.

### Imaging protocols

To obtain preoperative structural and diffusion-weighted images, we scanned all subjects at Baylor College of Medicine’s Core for Advanced Magnetic Resonance Imaging (CAMRI) in a Siemens MAGNETOM Prisma Fit 3T scanner (Erlangen, Germany) with a 32-channel head array coil. For the OCD cohort, we employed two different MRI acquisition protocols, with the specific protocol determined by the clinical trial design each subject participated in. Both protocols included a high resolution T1-weigheted magnetization-prepared rapid acquisition gradient-echo (MP-RAGE) sequences (256 sagittal slices, TR =2.4 s, and flip angle = 8°), with protocol-specific differences in echo time (TE = 2.41 ms vs 2.24 ms), spatial resolution (voxel size = 0.7 mm isotropic vs 0.8 mm isotropic), and field of view (FOV = 224x224x179 mm vs 256x282x205 mm). Also, a high resolution T2 variable-flip Turbo Spin Echo (TSE) sequence was acquired ( 208 sagittal slices, voxel size = 0.8 mm isotropic, TR = 3.2 s, TE = 563 ms, flip angle = 120°, and FOV = 240x256x166 mm, except one patient, who had 256 sagittal slices, 0.7-mm isotropic voxel size, and FOV = 224x224x179 mm). For the diffusion-weighted sequences, one protocol used single-shell, multi-band, high-angular-resolution with 268 axial volumes acquired with anterior-to-posterior phase encoding direction (including 12 b = 0 s/mm^2^ volumes and 256 diffusion-weighted directions at b = 1000 s/mm^2^), along with an additional 7 b = 0 s/mm^2^ axial volumes acquired with posterior-to-anterior phase encoding direction, voxel size = 2 mm isotropic, TR = 4.2 s, TE = 65 ms, flip angle = 90°, FOV = 256x256x128 mm, pixel bandwidth = 2300 Hz, and imaging frequency around 123.25 MHz. The other protocol used multi-shell, multi-band with 99 axial volumes (including 7 volumes at b = 0 s/mm^2^ and 46 diffusion-weighted directions acquired with opposite phase encoding directions at each of two b-values (b = 1250 s/mm^2^ and b = 2500 s/mm^2^), TR = 3.2 s, TE = 87 ms, flip angle = 90°, voxel size = 1.5 mm isotropic, FOV = 210x210x138 mm, pixel bandwidth = 1700 Hz, and imaging frequency = 123.25 MHz.

For the TRD cohort, the MRI scan sequences included a high resolution T1-weighted MP-RAGE sequence with 208 sagittal slices, TR = 2.4 s, TE = 2.24 ms, flip angle = 8°, voxel size = 0.8-mm isotropic, and FOV = 240x2562x166 mm; a high resolution T2 variable-flip TSE sequence with 208 sagittal slices, TR = 3.2 s, TE = 563 ms, flip angle = 120°, voxel size = 0.8 mm isotropic, and FOV = 240x256x166 mm; and a multi-shell, multi-band diffusion-weighted sequences with 99 axial volumes (including 7 volumes at b = 0 s/mm^2^ and 46 diffusion-weighted directions acquired with opposite phase encoding directions at each of b = 1000 s/mm^2^ and b=2000 s/mm^2^), TR = 3.4 s, TE = 85.8 ms, flip angle = 78°, voxel size = 1.5 mm isotropic, FOV = 210x210x138 mm, pixel bandwidth = 1700 Hz, and imaging frequency = 123.25 MHz.

We acquired intra-operative clinical CT scans following lead implantation on a Philips iCT 256 system, with 250-mm reconstruction diameter and contiguous slices of 0.67-mm thickness (0.67 mm spacing). The reconstructed images had an in-plane matrix size of 512 x 512 pixels, with a view size of 1664 x 1236 pixels.

### MRI preprocessing and tractography

We first pre-processed images using FreeSurfer version 7.3.2 (https://surfer.nmr.mgh.harvard.edu) to divide brain anatomical structures within each subject’s native space (Fischl et al., 2004). We then performed a visual inspection to enable necessary corrections to the automatic segmentations.

We preprocessed raw diffusion-weighted images using MRtrix3 (https://www.mrtrix.org) tools for denoising (Veraart et al., 2016) and Gibbs ring correction (Kellner et al., 2016). Next, we performed corrections for susceptibility-induced geometric distortions followed by motion and eddy current correction using Functional Magnetic Resonance Imaging for the Brain Software Library’s (FMRIB) Software Library (FSL’s) topup and eddy functions(Andersson et al., 2016). During this step, if diffusion-weighted images were acquired with opposite phase encoding directions, volumes with the same diffusion sensitization were paired and combined to estimate the susceptibility distortion field. If only b = 0 images were available with opposite phase encoding directions, the paired b = 0 images were used for distortion field estimation. Subsequently, we performed bias field correction using Advanced Normalization Tools (ANTs) (Tustison et al., 2010).

In addition, we co-registered the skull-stripped b0 image, T2-weighted image, and T1-weighted images using FMRIB’s Linear Image Registration Tool (FLIRT) affine transformation matrix, to generate a Freesurfer-derived five-tissue type (5TT) segmentation (i.e., cortical gray matter, subcortical gray matter, white matter, cerebrospinal fluid, and pathological tissue) aligned to the diffusion-weighted images. Skull stripping was conducted using the FSL’s Brain Extraction Tool (BET) (Smith, 2002) or the HD-BET (Isensee et al., 2019). As in (Sretavan et al., 2024), the 5TT segmentation was modified such that the caudate nucleus, putamen, and nucleus accumbens were reassigned from the subcortical gray matter compartment to the white matter compartment because many ventral PFC-ALIC fibers pass through fascicles embedded in these subcortical structures.

Moreover, we applied Constrained Spherical Deconvolution (CSD) to estimate fiber orientation distribution separately for gray matter, white matter, and cerebrospinal fluid (Tournier et al., 2004). From there, we used predefined regions-of-interest (ROIs) to generate streamlines. We extracted the ALIC mask image from the Johns Hopkins University ICBM-DTI-81 White Matter Labeled Atlas (Mori et al., 2008) and a thalamic mask from the Harvard-Oxford cortical and subcortical structural atlas. As previously described by Banks et al,2022, we thresholded the thalamic mask at 0.5, further refined to exclude voxels included within 1 voxel distance of the ALIC mask (Banks et al., 2022). The brainstem mask was drawn manually on 6 slices of the MNI152 1-mm standard brain inferior to the subthalamic nucleus, also as described previously (Banks et al., 2022). Following (Sretavan et al., 2024), we subdivided the PFC into eleven subregions per hemisphere using the FreeSurfer’s Desikan-Killiany Atlas (Desikan et al., 2006). These ROIs were the caudal middle frontal cortex, rostral middle frontal cortex, superior frontal cortex, caudal anterior cingulate cortex (cACC), rostral anterior cingulate cortex (rACC) (with the rostral cingulate subdivided into dorsal and ventral rostral anterior cingulate), pars opercularis, pars triangularis, par orbitalis, central OFC, and mOFC. The rACC was further subdivided at the level of the SCC (z = 76 mm) in MNI space that was then registered to subject native space due to the unique trajectory the ventral fibers follow which differ from the dorsal fibers (Sretavan et al., 2024). We registered all ROIs to each subject diffusion-weighted image by applying ANTs-derived transformation matrices to warp the ROIs from MNI152 template or the subject T1-weighted image/FreeSurfer labels into native diffusion (b = 0) space.

Finally, we performed probabilistic tractography using Anatomy Constrained Tractography (ACT) (Smith et al., 2015) in MRtrix3 within the ALIC mask. Using the “tckgen” function, we generated one million streamlines passing through the ALIC (with a maximum angle of 22.5 degrees and a maximum length of 250 mm), and then we used “tckedit” function to select only streamlines passing through the ALIC mask and terminating in the other ROIs (PFC subregions, thalamus, and brainstem).

### ALIC topography

To statistically assess the topography of PFC fibers traversing the ALIC along the dorsoventral axis, we quantified the dorsoventral coordinates of streamline-weighted centroids for each subject at the level of the decussation of the anterior commissure (in OCD patients only, as there were only 5 TRD patients). We then grouped these values into dorsally, centrally, and ventrally projecting PFC regions in each hemisphere based on a priori hypotheses (Safadi et al., 2018; Sretavan et al., 2024) and then performed Wilcoxon signed-rank test with Bonferroni correction adjusted for multiple comparisons to evaluate cortical topographies. The dorsal PFC group included the caudal middle frontal, rostral middle frontal and superior frontal regions; the central PFC group included the cACC, dorsal rACC, pars opercularis, pars triangularis, and pars orbitalis regions; and the ventral PFC group included lOFC, mOFC, and ventral rACC regions. For the subcortical regions, we quantified the centroids along the mediolateral axis and then performed Wilcoxon signed-rank test to evaluate subcortical topographies.

### sEEG Electrode Localization

Electrodes were localized using the software pipeline intracranial Electrode Visualization (iELVis; (Groppe et al., 2017)) and plotted across patients on an average brain using Reproducible Analysis & Visualization of iEEG (RAVE; (Magnotti et al., 2020)). For each patient, DICOM images of the preoperative T1 anatomical MRI and the postoperative Stealth CT scans were acquired and converted to NIfTI format (Li et al., 2016). Functional Magnetic Resonance Imaging for the Brain Software Library’s (FMRIB’s) Linear Image Registration Tool (FLIRT) was used to align the preoperative MRI scan and the postoperative CT scan. The output co-registered CT image was visualized in BioImage Suite (version 3.5β1; (Joshi et al., 2011)) to determine electrode coordinates. Then iELVis was used to determine the coordinates in patient native space (Yang et al., 2012). The orbitofrontal electrode coordinates were converted into average MNI152 space and then visualized using RAVE (Magnotti et al., 2020). The vPFC electrode coordinates were plotted together on an average glass brain and colored by patient.

### Single-pulse electrophysiological stimulation and recording

Monopolar cathodic single pulse stimulation was delivered via a Blackrock CereStim R96 stimulator (Blackrock Microsystems, Utah, USA) (Adkinson et al., 2022; Sheth et al., 2022; Schmid et al., 2024). Stimulation was delivered using biphasic pulses with amplitudes an amplitude of 5 mA, a pulse width of 180 µs with a 100 µs interphase gap. For each subject and at each stimulated contact, a total of 315 stimulation pulses were used, delivered every 600ms with a random jitter ranging from 1 to 200 ms. Neural signals from sEEG probes located in the vPFC (vmPFC, OFC, vlPFC) were recorded at 30 kHz sampling rate with a 1^st^ order high-pass Butterworth filter at 0.3 Hz on a 256-channel Blackrock Cerebus system (Blackrock Microsystems, UT, USA).

### Single-pulse evoked potential preprocessing and analysis

Custom MATLAB scripts (Mathworks, MA, USA) were used for all signal preprocessing and analysis. First, we visually inspected the sEEG signals for line noise, excessive recording artifact and jaw contraction contamination, leading to the exclusion of 00-04 channels (min-max range across our sample). The neural signal around each single pulse onset was identified, and a window of interest was defined (150 ms preceding and 300 ms following each pulse). The average waveform for each sEEG contact (across the 315 stimulation pulses) was z-scored with respect to baseline (120-20 ms preceding the pulse) and peaks exceeding 6 z-scores in the 10-100 ms window following stimulation were detected using the MATLAB function *findpeaks*. The prominence of the largest peak (if any) was used to quantify the evoked potential.

### Streamline analysis for TRD patients

To determine structural connectivity between DBS stimulating contacts and vPFC sEEG recording contacts, we quantified the number of streamlines connecting each contact pair. For each vPFC contact, we performed probabilistic tractography using ACT in MRtrix3, with a 3 mm radius spherical seed mask centered on the contact. Using the “tckgen” function, we generated 500,000 streamlines from each vPFC seed and then applied “tckedit” function to select only tracts traversing the ALIC. From these filtered tracts, streamlines passing through each VC/VS DBS contact were extracted using a 1.5 mm radius spherical ROI mask centered on each DBS contact location again using “tckedit”.

To examine whether white matter structural connectivity (streamline count) is associated with evoked potential amplitude, we first performed a global Spearman rank correlation across all contact pairs pooled from all subjects. To account for the non-independence of contact pairs within subjects, we performed linear mixed-effect models with log-transformed streamline count as a fixed effect of CCEP amplitude and subject identity as a random intercept, thereby capturing between-subject differences in baseline amplitude.

## Results

### PFC-ALIC topography replicated using diffusion tractography in treatment-resistant OCD patients

We applied a variant of our anatomically validated diffusion tractography pipeline (Sretavan et al., 2024) to data from 18 treatment-resistant OCD patients to visualize the spatial distribution of tracts from PFC regions through the ALIC (Figure 1). In this cohort, the ALIC demonstrated consistent topography of PFC fibers that matched underlying anatomical principles (Figure 1). Specifically, at the level of the decussation of the anterior commissure and by comparing dorsal to ventral regions, centroids of tracts originating from the dlPFC (rostral and caudal middle frontal cortices) and the dmPFC (superior frontal cortex) were located more dorsally within the ALIC than centroids of tracts originating from the ventral PFC (central OFC, mOFC, and ventral rACC) (Left W=171, Z=3.72, p < 0.001; Right W=153, Z=3.62, p < 0.001). Comparing dorsal to central regions showed that fibers from the vlPFC (including pars opercularis, pars triangularis, and pars orbitalis) and from the dorsal ACC (dorsal rACC and cACC) were located in the center of the ALIC (Left W=171, Z=3.72, p < 0.001; Right W=272, Z=3.72, p < 0.001) (Figure 1). Fibers from the ventral PFC (mOFC, central OFC, and ventral rACC) were positioned most ventrally (Left W = 171, Z = 3.72, p < 0.001; Right W = 168, Z = 3.59, p < 0.001) when comparing central to ventral areas. These results confirm that the known topographic organization of PFC fibers through the ALIC, occupying stereotyped dorsoventral positions, is preserved in a cohort of treatment-resistant OCD patients. As we planned to evaluate their PFC-ALIC CCEPs in the TRD cohort, we also applied the tractography pipeline to the TRD patients (n=5). Although too few in number to test statistically, visual inspection revealed similar topography.

**Figure 1.**
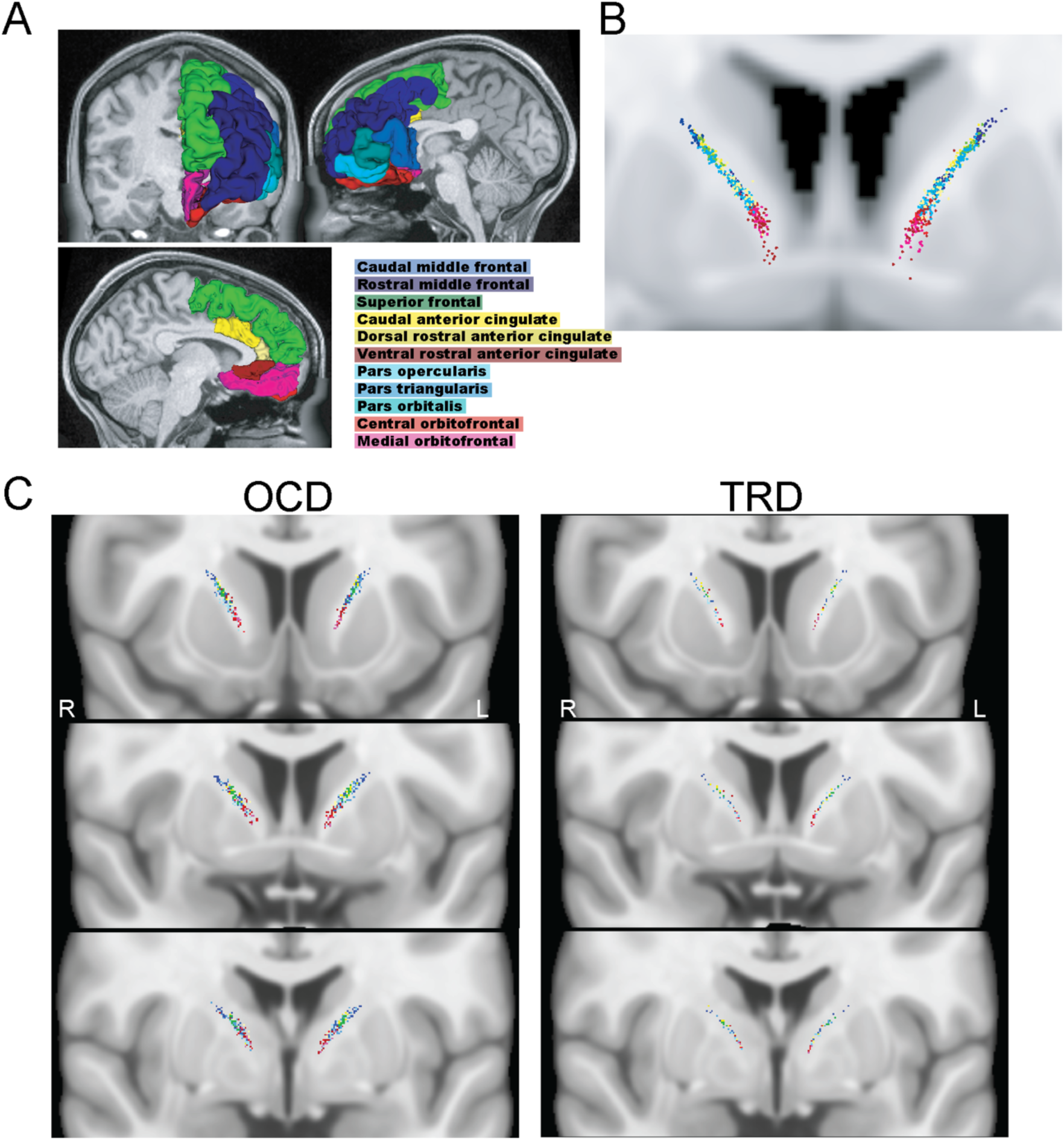
PFC topography in the ALIC of treatment-resistant psychiatric patients. A. PFC divisions used to generate segmented ALIC tracts, as described in Sretavan et al., 2024. B. PFC-ALIC tract locations (centroids) match known topography with stereotyped dorsoventral organization in healthy controls, from Sretavan et al., 2024. C. PFC-ALIC tract locations (centroids) match known topography with stereotyped dorsoventral organization in treatment-resistant OCD and TRD patients.

### Subcortical topographical organization in the ALIC

In prior work (Banks et al., 2022), we demonstrated that ALIC fibers targeting the thalamus are located medially to those targeting the brainstem in a cohort of healthy control patients, a finding that replicates prior microscopy work (Axer et al., 1999). Here, we replicated this topographic organization in the treatment-resistant OCD cohort, and by comparing the centroids along the mediolateral axis of the thalamus and the brainstem in both hemispheres, results showed that the thalamus was located medially compared to brainstem with a significant result only in the left hemisphere ( Left W=142, p<0.01, Right W=40, p=0.08).

### vPFC-ALIC topography in CCEPs

In the TRD cohort, we stimulated individual contacts of ALIC DBS electrodes while recording from the ipsilateral vPFC (Figure 3A-B). Prior work, as well as the tractography results above, demonstrate a medial-lateral vPFC to ventral-dorsal VC projection transformation (Lehman et al., 2011; Sretavan et al., 2024). In order to account for interindividual variability in electrode placement that caused alternative topographies, we normalized CCEP amplitudes within each subject and then utilized a linear regression analysis across all subjects which demonstrated a significant spatial relationship between CCEP amplitude and the mediolateral coordinates of recording electrodes during stimulation along the dorsoventral ALIC axis in the left hemisphere (β = −0.0026, p = 0.049), but not the right hemisphere (β = 0.00118, p = 0.49). To visualize this effect, we plotted whether stronger CCEP amplitudes were associated with more medial vPFC sEEG contacts, or more lateral vPFC sEEG contacts. In keeping with prior anatomical and tractography literature, in both hemispheres, positive correlations (yellow spheres in Figure 3C-D, corresponding to increased CCEP amplitudes at more medial vPFC contacts) were present in the ventral ALIC, whereas negative correlations (blue spheres in Figure 3C-D, corresponding to increased CCEP amplitudes at more lateral vPFC contacts) were present in more dorsal ALIC sites. This spatial organization highlights an effective connectivity pattern linking ventral ALIC stimulation sites to medial vPFC responses, and dorsal ALIC stimulation sites to lateral vPFC responses.

**Figure 2.**
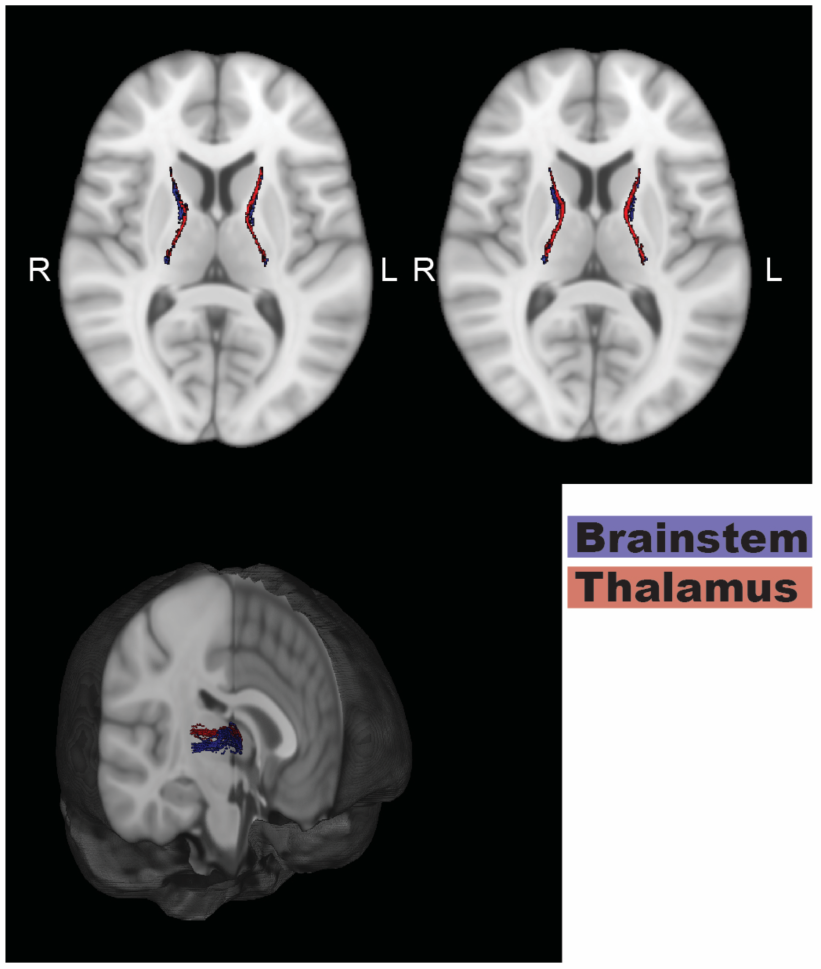
Thalamus and brainstem topography in the ALIC of treatment-resistant OCD patients. Thalamus/brainstem-ALIC tract locations (centroids) match known topography of a medial-lateral distribution, as established by anatomy and tractography in healthy controls.

**Figure 3:**
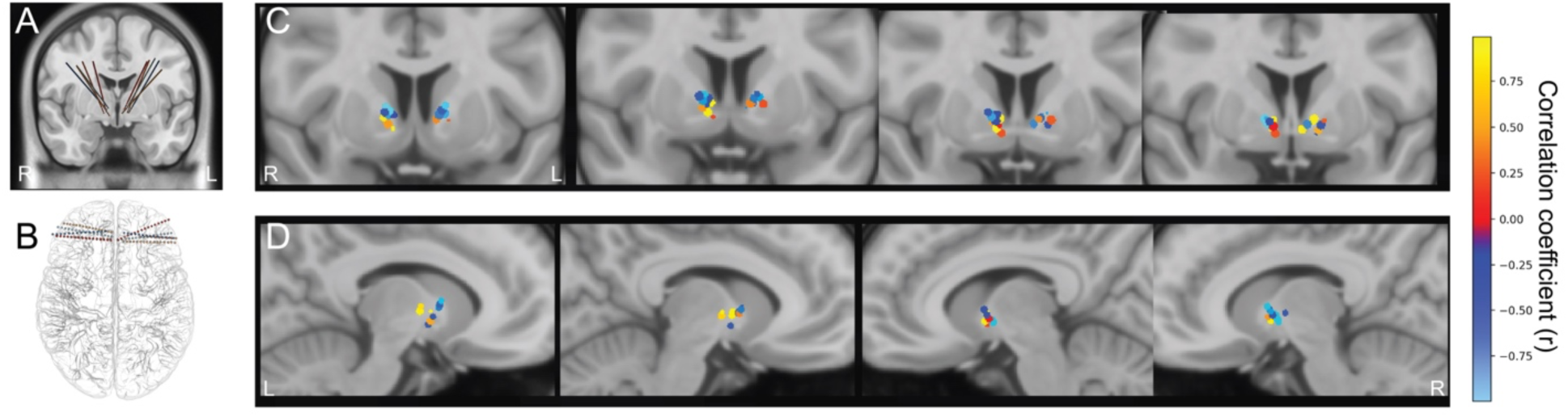
ALIC-vPFC CCEP topography. **A.** ALIC DBS electrode placement in TRD patients. Individual ALIC contacts were stimulated to generate evoked potentials in the vPFC. **B.** sEEG leads in the vPFC of TRD patients. sEEG electrodes sampled the medial-lateral extent of the vPFC, with the exact placement varying by patient and hemisphere. **C-D.** Each DBS ALIC contact is shown in MNI space, colored according to the correlation between vPFC sEEG contact medial-lateral position and CCEP amplitude. Hotter colors indicate greater response in medial vPFC; cooler colors indicate greater responses in lateral vPFC. Overall, in the left hemisphere, ventral ALIC stimulation sites were associated with greater CCEP responses in the medial vPFC, whereas dorsal ALIC stimulation sites were associated with greater CCEP responses in the lateral vPFC, recapitulating established topographies.

### White Matter Connectivity and CCEP amplitude

To evaluate whether the structural connectivity pattern between the ALIC stimulating contacts and vPFC recording contacts mirrors the functional connectivity pattern identified from CCEPs, we tested the correlation between contact pairs’ streamline counts and CCEP amplitudes. Global Spearman rank correlation revealed a non-significant positive correlation in the left hemisphere (ρ = 0.177, p = 0.107) and a non-significant negative association in the right hemisphere (ρ = -0.085, p = 0.358). The linear mixed-effects model also failed to show any significant results (Left β =0.248, p = 0.224, Right β = -0.749, p = 0.453). The random intercept variance indicated that between-subject differences in baseline CCEP amplitude accounted for approximately 60% of total variance in the left hemisphere (Group Var = 41.625, residual = 27.571) and 50% of total variance in the right hemisphere (Group Var = 499.18, residual = 474.84), suggesting that subject-level effects dominate the variance structure and limit the statistical power to detect a fixed within-subject streamline–CCEP effect. Visual inspection of ALIC DBS contact locations and ventral PFC region centroids revealed inter-subject variability in electrode placement relative to the white matter pathways connecting these regions (Figure 4). In both hemispheres, but particularly on the right, centroids were mostly located ventral to contact locations.

**Figure 4:**
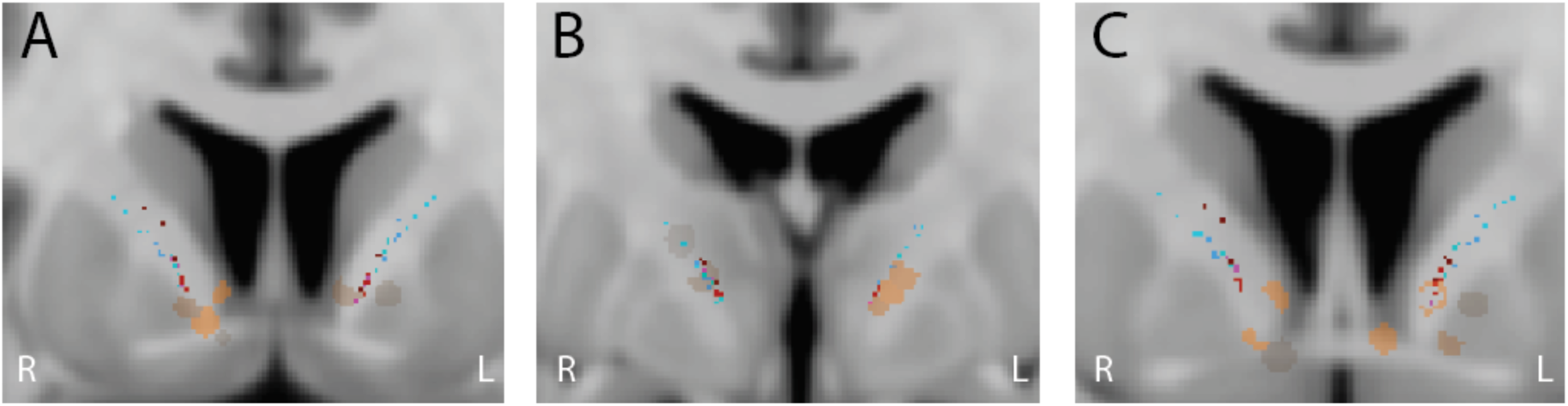
vPFC centroids topography relative to the ALIC stimulating DBS contacts. **A-C.** TRD Patients ALIC DBS electrode placement (orange-brown color) relative to vPFC centroids.

## Discussion

In this study, we investigated the topographical organization of prefrontal and subcortical fibers traversing the ALIC, a major conduit for limbic and cognitive pathways, and an established DBS target for treatment-resistant OCD. Because dysregulation within and across prefrontal circuits is central to OCD pathology (Ahmari and Rauch, 2022), characterizing the spatial distribution of these pathways provides an essential framework for understanding the circuit-level effects of ALIC stimulation. We were able to analyze diffusion tractography collected from treatment-refractory psychiatric patients as well as a highly unique dataset of CCEPs collected from TRD patients.

Our diffusion tractography results indicate that the dorsoventral topography of PFC projections in the ALIC is preserved in patients with treatment-refractory OCD, consistent with recent structural mapping studies in healthy individuals (Sretavan et al., 2024) and prior tract-tracing work in nonhuman primates (Lehman et al., 2011; Safadi et al., 2018). Furthermore, previously described medial-lateral organization of the ALIC, with brainstem fibers positioned laterally to thalamus fibers (Banks et al., 2022; Axer et al., 1999), could also be reconstructed in treatment-resistant OCD patients. Thus, large-scale organizational principles remain intact despite the functional disturbances associated with the disorder.

To assess functional dynamics, we analyzed CCEP responses evoked in sEEG electrodes by single-pulse stimulation via ALIC DBS electrodes in TRD patients. CCEP amplitudes varied systematically with both the dorsoventral position of the stimulating electrode and the mediolateral position of the orbitofrontal recording contact. Specifically, stimulation delivered from more ventral DBS electrodes produced stronger responses in medial vPFC regions, indicating a ventral->medial / dorsal->lateral functional coupling pattern that was prominent in the left hemisphere. This directional relationship aligns with anatomical topographies described in human diffusion MRI studies and primate tract-tracing that showed ventral ALIC pathways preferentially containing axons from medial OFC and ventromedial prefrontal areas, whereas dorsal ALIC fibers connect to lateral OFC and vlPFC (Jbabdi et al., 2013; Safadi et al., 2018; Sretavan et al., 2024). This anatomical pattern is partially the result of different entry points to the ALIC: medial vPFC fibers traverse the nucleus accumbens to reach the ALIC from its ventral border, while lateral vPFC fibers circumvolve the putamen to reach the ALIC from its dorsal border. Our findings suggest that the intrinsic ventrodorsal–mediolateral organization of frontal ALIC fibers is reflected in effective connectivity measured directly in humans.

CCEPs are typically obtained during epilepsy monitoring, where they help reveal the network dynamics associated with the epileptogenic zone (Zhao et al., 2019; Qiang et al., 2025). Outside of epilepsy monitoring, CCEPs are not routinely obtained. However, there is increasing interest in how they might be used to evaluate network integrity and engagement during neuromodulation treatment (Waters et al., 2018; Sheth et al., 2022). Understanding how they relate to the underlying principles of anatomical connectivity will be essential for maximizing the use of CCEPs in neuromodulation and epilepsy.

Although CCEPs provide direct measures of network-level effective connectivity, they do not specify the anatomical pathways that carry the signal, nor do they fully account for variability in latency and amplitude across recording sites (Silverstein et al., 2020). Conversely, diffusion tractography is the primary non-invasive tool that offers detailed structural connectivity maps, but it is limited by false-positives and anatomically implausible bundles generated by tractograms (Maier-Hein et al., 2017; Grier et al., 2020). To overcome the limitations of each method, multimodal approaches are needed. We used information from tract-tracing in nonhuman primates and diffusion tractography in healthy control subjects to generate anatomically plausible pipelines (Lehman et al., 2011; Safadi et al., 2018; Sretavan et al., 2024).

The OFC is a central hub for emotional regulation and reward-based decision-making, but it is highly heterogeneous, with both medial-lateral and anterior-posterior gradients in structure and function (Rushworth et al., 2011). The medial OFC, along with the ventromedial prefrontal cortex, is associated with reward, value signaling, and decision making (Plassmann et al., 2007; Boorman et al., 2009; Noonan et al., 2010). More lateral aspects of OFC seem to be more strongly implicated in learning (Noonan et al., 2010). Medial OFC functionally connects with limbic and autonomic regions, while lateral OFC functionally connects with cognition and language regions (Zald et al., 2014). These distinctions are also present in psychiatric disorders. In depression, the lateral OFC is hyper-connected and highly sensitized, whereas the medial OFC shows reduced connectivity as well as reduced reward responses (Zhang et al., 2024). In OCD, the medial OFC is hyper-connected, and its excessive coupling with basal ganglia circuits is thought to contribute directly to symptom severity (Beucke et al., 2013). These different alterations in the OFC highlight its shared and unique vulnerabilities across psychiatric disorders. One possibility is that, in DBS and capsulotomy treatments, medial vs lateral OFC fiber engagement produces different outcomes, mediated by targeting different portions of the ALIC along the dorsoventral axis.

There are multiple specific psychiatric DBS targets involving the ALIC, including the BNST (Mosley et al., 2021; Meyer et al., 2024), VS (Greenberg et al., 2006), nucleus accumbens (Sturm et al., 2003; Denys et al., 2010), superolateral branch of the medial forebrain bundle (Schlaepfer et al., 2013; Coenen et al., 2017), and STN (Schneider et al., 2003). Because of the topographic organization of the ALIC, all of these sites have the potential to have an impact on specific PFC regions, depending on the precise location stimulated. One proposal is that the most impactful sites for these interventions are the limbic OFC and ACC pathways that are critical for motivation (Haber et al., 2021). There is additional precision offered by the topography of OFC fibers within the ALIC, although which of those fibers should be stimulated for optimal outcomes remains unknown.

Intersubject variability and reproducibility of diffusion tensor imaging are important considerations in diffusion tractography. Previous work has shown that diffusion metrics and tractography measures are generally reproducible within subjects, whereas differences between subjects (intersubject variability) across different brain regions are substantial (Heiervang et al., 2006). Such variability may arise from scanner-related factors, including b0 field inhomogeneities, gradient stability, signal-to-noise fluctuations, and software upgrades, as well as subject-related effects such as head motion and positioning within the scanner field of view (Veenith et al., 2013). Despite individual variability in the precise trajectories of these fibers (Makris et al., 2016; Safadi et al., 2018), their broader topographic organization remains remarkably consistent, enabling fiber-informed targeting strategies. Incorporating patient-specific tractography may therefore help optimize electrode placement and enhance therapeutic outcomes by aligning stimulation with circuits most relevant to symptom improvement.

Our study has several limitations. First, the sample size of our TRD cohort was small (n=5), which limited statistical power for the CCEP analysis. Second, our data acquisition protocols were neither standardized nor aligned with those used in our prior work on healthy controls (Banks et al., 2022; Sretavan et al., 2024). This precluded a direct comparison of healthy controls and patient populations. Still, despite data acquisition differences, we were able to successfully implement the previously established analysis pipelines and reproduce known topographies. This highlights the method’s robustness to acquisition-related variability and supports its potential suitability for broader use in multi-site studies. Finally, although additional prefrontal (non-vPFC) sEEG electrodes were implanted in TRD patients (Sheth et al., 2022), our analysis was restricted to vPFC electrodes placed along the mediolateral axis. Electrodes placed in other regions traversed multiple anatomical gradients, and thus were excluded, as their trajectories were difficult to reliably map along a single spatial axis. Future studies with larger cohorts, and more diverse electrode trajectories will be critical to validate these findings and to extend this framework toward more comprehensive, patient-specific circuit mapping for therapeutic targeting.

We failed to observe a significant correlation between ALIC-vPFC CCEP response amplitude and structural connectivity; such a correlation has been observed in other, larger datasets (Crocker et al., 2021; Paulk et al., 2022; Schmid et al., 2024; Danstrom et al., 2026). Moreover, in the left but not the right hemisphere, we did observe a relationship between medial-lateral vPFC recording location, dorsal-ventral ALIC stimulation location, and CCEP amplitude. Although limited by a small number of observations, there are several other possible explanations for our null results. One is that medial vPFC-ALIC fibers are more difficult to track through the nucleus accumbens and anterior commissure (Jbabdi et al., 2013); although our medial vPFC streamline centroids are rightfully positioned ventral in the ALIC relative to lateral vPFC centroids, it is possible that the placement is still not ventral enough, leading to a disconnect between the true anatomy and the diffusion tractography. In this case, CCEPs patterns may more accurately reflect the underlying anatomy than diffusion tractography does. Another possibility is that, particularly in the right hemisphere, there are simply few vPFC fibers near the ALIC contacts.

## Conclusion

This study demonstrates that the topographic organization of prefrontal projections within the ALIC, as well as subcortical projections from the thalamus and brainstem, is preserved in treatment-resistant OCD patients, consistent with findings in healthy individuals. Using both diffusion tractography and CCEP recordings, our findings link known fiber topographies to stimulation–response patterns in the vPFC, with a lateralized ventromedial coupling in the left hemisphere. Together, these results underscore the importance of mapping white matter functional and structural topography for circuit-informed DBS targeting, with implications for refining DBS therapy and advancing more precise, patient-specific neuromodulation strategies.

## Acknowledgments

The authors acknowledge funding for this study from NIH UH3NS103549, R01MH1340597, UH3NS100549, and the Robert and Janice McNair Foundation. We thank Victoria Gates, Josh Adkinson, and Denise Oswalt for assistance with data collection.

---

**Supplemental Table 1:** MNI152 coordinates of lead DBS electrodes in OCD patients.

| Patient ID | VC/VS<br>Contacts | Left Hemisphere |  |  | Right Hemisphere |  |  |
| --- | --- | --- | --- | --- | --- | --- | --- |
|  |  | x (mm) | y (mm) | z (mm) | x (mm) | y (mm) | z (mm) |
| OCD001 | Contact 1 | -10.47 | 2.86 | -9.36 | 11.62 | 5.14 | -6.25 |
|  | Contact 2 | -11.79 | 4.69 | -6.86 | 13.28 | 6.81 | -3.61 |
|  | Contact 3 | -13.17 | 6.54 | -4.33 | 15.06 | 8.52 | -1.12 |
|  | Contact 4 | -14.61 | 8.35 | -1.87 | 16.82 | 10.15 | 1.14 |
| OCD002 | Contact 1 | -9.83 | 0.41 | -4.42 | 8.80 | 2.79 | -8.48 |
|  | Contact 2 | -11.80 | 2.106 | -1.97 | 10.62 | 4.56 | -5.83 |
|  | Contact 3 | -13.69 | 3.68 | 0.403 | 12.41 | 6.24 | -3.23 |
|  | Contact 4 | -15.54 | 5.19 | 2.72 | 14.25 | 7.94 | -0.72 |
| OCD003 | Contact 1 | -11.54 | 7.05 | -7.61 | 9.96 | -0.35 | -4.62 |
|  | Contact 2 | -13.16 | 8.31 | -4.61 | 11.45 | 1.49 | -2.43 |
|  | Contact 3 | -14.91 | 8.93 | -1.57 | 12.94 | 2.75 | 0.13 |
|  | Contact 4 | -16.81 | 8.98 | 1.55 | 14.56 | 3.51 | 3.18 |
| OCD004 | Contact 1 | -6.22 | 0.65 | -2.94 | 9.006 | 1.47 | -2.82 |
|  | Contact 2 | -8.22 | 2.49 | -0.84 | 10.95 | 3 | -0.43 |
|  | Contact 3 | -10.28 | 3.87 | 1.65 | 12.99 | 3.97 | 2.27 |
|  | Contact 4 | -12.45 | 4.64 | 4.58 | 15.05 | 4.35 | 5.42 |
| OCD005 | Contact 1 | -3.15 | -2.06 | -8.57 | 6.16 | -0.48 | -8.97 |
|  | Contact 2 | -5.53 | -1.04 | -6.45 | 8.29 | 0.07 | -6.18 |
|  | Contact 3 | -7.81 | 0.03 | -4.32 | 10.36 | 0.709 | -3.52 |
|  | Contact 4 | -9.96 | 1.12 | -2.18 | 12.43 | 1.38 | -0.97 |
| OCD006 | Contact 1 | -5.51 | -1.34 | -8.207 | 5.61 | 0.23 | -7.07 |
|  | Contact 2 | -7.83 | 0.78 | -6.55 | 7.57 | 2.27 | -4.87 |
|  | Contact 3 | -10.01 | 2.49 | -4.57 | 9.53 | 3.79 | -2.49 |
|  | Contact 4 | -12.18 | 3.92 | -2.03 | 11.64 | 4.81 | 0.21 |
| OCD007 | Contact 1 | -7.52 | -0.8 | -5.27 | 9.29 | 0.59 | -5.22 |
|  | Contact 2 | -9.44 | 1.10 | -3.6 | 11.01 | 2.53 | -3.49 |
|  | Contact 3 | -11.36 | 2.44 | -1.5 | 12.83 | 3.88 | -1.37 |
|  | Contact 4 | -13.44 | 3.23 | 1.02 | 14.85 | 4.69 | 1.15 |
| OCD008 | Contact 1 | -11.13 | 2.93 | -5.62 | 8.92 | 3.74 | -5.87 |
|  | Contact 2 | -12.51 | 4.35 | -3.24 | 10.76 | 5.46 | -3.90 |
|  | Contact 3 | -13.9 | 5.77 | -0.78 | 12.61 | 7.19 | -1.87 |
|  | Contact 4 | -15.31 | 7.19 | 1.73 | 14.51 | 8.88 | 0.17 |
| OCD009 | Contact 1 | -7.33 | -2.78 | -5.86 | 9.84 | -1.28 | -5.14 |
|  | Contact 2 | -8.94 | -0.60 | -4.13 | 11.38 | 0.39 | -2.76 |
|  | Contact 3 | -10.63 | 1.04 | -1.91 | 13.06 | 1.35 | -0.11 |
|  | Contact 4 | -12.43 | 2.13 | 0.85 | 14.94 | 1.58 | 2.78 |
| OCD010 | Contact 1 | -11.32 | 3.56 | -5.79 | 8.36 | 3.19 | -4.53 |
|  | Contact 2 | -13.11 | 4.37 | -3.11 | 10.14 | 4.03 | -1.97 |
|  | Contact 3 | -15.00 | 5.09 | -0.35 | 12.06 | 4.72 | 0.61 |
|  | Contact 4 | -16.92 | 5.75 | 2.38 | 14.01 | 5.32 | 3.18 |
| OCD011 | Contact 1 | -7.32 | -0.11 | -3.42 | 8.67 | 2.35 | -4.06 |
|  | Contact 2 | -9.28 | 1.67 | -1.21 | 10.45 | 4.40 | -1.98 |
|  | Contact 3 | -11.13 | 3.44 | 1.04 | 12.27 | 6.44 | 0.08 |
|  | Contact 4 | -12.97 | 5.15 | 3.29 | 14.08 | 8.52 | 2.13 |
| OCD012 | Contact 1 | -8.38 | 5.06 | -5.03 | 9.07 | 3.73 | -6.69 |
|  | Contact 2 | -10.13 | 5.86 | -2.48 | 10.93 | 4.52 | -4.17 |
|  | Contact 3 | -11.83 | 6.62 | 0.01 | 12.72 | 5.27 | -1.63 |
|  | Contact 4 | -13.52 | 7.3 | 2.44 | 14.54 | 5.91 | 0.91 |
| OCD013 | Contact 1 | -10.23 | 5.06 | -9.03 | 10.31 | 5.65 | -8.58 |
|  | Contact 2 | -12.01 | 6.37 | -6.78 | 11.91 | 7.18 | -6.41 |
|  | Contact 3 | -13.79 | 7.65 | -4.48 | 13.44 | 8.702 | -4.2 |
|  | Contact 4 | -15.61 | 8.91 | -2.15 | 14.95 | 10.21 | -1.94 |
| OCD014 | Contact 1 | -10.97 | 0.38 | -4.61 | 11.03 | 1.21 | -3.51 |
|  | Contact 2 | -12.39 | 1.67 | -1.98 | 12.59 | 3.00 | -1.31 |
|  | Contact 3 | -13.75 | 2.89 | 0.67 | 14.22 | 4.77 | 0.92 |
|  | Contact 4 | -15.16 | 4.01 | 3.36 | 15.96 | 6.52 | 3.13 |
| OCD015 | Contact 1 | -11.17 | 5.31 | -8.60 | 6.31 | 1.10 | -8.22 |
|  | Contact 2 | -12.61 | 6.42 | -5.73 | 8.1 | 2.27 | -5.69 |
|  | Contact 3 | -13.97 | 7.57 | -2.97 | 9.86 | 3.51 | -3.26 |
|  | Contact 4 | -15.31 | 8.72 | -0.25 | 11.64 | 4.80 | -0.90 |
| OCD016 | Contact 1 | -8.94 | -0.69 | -2.72 | 5.58 | -0.07 | -5.3 |
|  | Contact 2 | -11.05 | 1.42 | -0.67 | 8.17 | 1.98 | -2.93 |
|  | Contact 3 | -13.07 | 2.89 | 1.72 | 10.6 | 3.28 | -0.55 |
|  | Contact 4 | -14.97 | 3.75 | 4.51 | 13.09 | 3.8 | 1.97 |
| OCD017 | Contact 1 | 6.4 | -0.96 | -5.1 | 6.4 | -0.96 | -5.1 |
|  | Contact 2 | 8.05 | 0.76 | -3.25 | 8.05 | 0.76 | -3.25 |
|  | Contact 3 | 9.71 | 1.97 | -1.01 | 9.71 | 1.97 | -1.01 |
|  | Contact 4 | 11.43 | 2.55 | 1.53 | 11.4 | 2.55 | 1.53 |
| OCD018 | Contact 1 | -9.08 | -0.69 | -4.62 | 6.46 | -0.78 | -5.08 |
|  | Contact 2 | -10.9 | 1.41 | -2.97 | 8.33 | 1.65 | -3.87 |
|  | Contact 3 | -12.71 | 2.98 | -0.88 | 10.06 | 3.67 | -2.11 |
|  | Contact 4 | -14.61 | 3.96 | 1.65 | 11.84 | 5.24 | 0.18 |

**Supplemental Table 2:** MNI152 coordinates of lead DBS electrodes in TRD patients.

| Patient ID | VC/VS<br>Contacts | Left Hemisphere |  |  | Right Hemisphere |  |  |
| --- | --- | --- | --- | --- | --- | --- | --- |
|  |  | x (mm) | y (mm) | z (mm) | x (mm) | y (mm) | z (mm) |
| TRD001 | Contact 1 | -6.21 | -0.85 | -3.14 | 9.27 | -0.67 | -3.76 |
|  | Contact 2 | -7.45 | -0.29 | -1.77 | 10.09 | -0.08 | -2.25 |
|  | Contact 3 | -8.63 | 0.29 | -0.34 | 11.18 | 0.56 | -0.48 |
|  | Contact 4 | -9.71 | 0.85 | 1.21 | 12.11 | 1.15 | 1.14 |
| TRD002 | Contact 1 | -10.29 | -3.85 | -1.11 | 11.36 | -4.22 | 0.4 |
|  | Contact 2 | -11.6 | -3.64 | 1.16 | 12.62 | -3.93 | 2.62 |
|  | Contact 3 | -13.19 | -3.54 | 3.46 | 13.82 | -3.61 | 4.82 |
|  | Contact 4 | -14.77 | -3.25 | 5.71 | 15.3 | -3.23 | 7.3 |
| TRD003 | Contact 1 | -4.39 | 0.8 | -3.52 | 7.34 | 2.23 | -5.33 |
|  | Contact 2 | -6.68 | 2.09 | -1.27 | 9.51 | 3.26 | -3.19 |
|  | Contact 3 | -9.39 | 3.59 | 1.11 | 11.68 | 4.62 | -1.08 |
|  | Contact 4 | -12.26 | 4.85 | 3.26 | 13.54 | 6.25 | 1.35 |
| TRD004 | Contact 1 | -8.81 | 4.13 | 0.76 | 8.04 | 2.81 | 0.88 |
|  | Contact 2 | -10.09 | 4.92 | 2.7 | 9.45 | 3.87 | 2.65 |
|  | Contact 3 | -11.36 | 5.48 | 4.62 | 11.17 | 4.92 | 4.19 |
|  | Contact 4 | -12.64 | 6.1 | 6.55 | 12.74 | 5.98 | 5.85 |
| TRD005 | Contact 1 | -9.41 | -0.72 | -4.31 | 9.21 | 1.43 | -1.49 |
|  | Contact 2 | -10.46 | 0.75 | -3.47 | 10.38 | 1.98 | 0.87 |
|  | Contact 3 | -11.59 | 1.42 | -1.11 | 12.79 | 3.57 | 2.88 |
|  | Contact 4 | -13.78 | 3.25 | 0.91 | 13.95 | 3.82 | 4.05 |

